# Bacteriophage susceptibility varies within chronic *Pseudomonas aeruginosa* lung infections

**DOI:** 10.64898/2026.10.13.758950

**Authors:** Noah S. B. Houpt, Jelly Vanderwoude, Stephen P. Diggle, Paul E. Turner

## Abstract

Phage therapy is receiving renewed interest in the face of rising rates of antibiotic resistance. One drawback of phages as antibiotic agents is their high host specificity: phages can often only kill a subset of strains within a bacteria species. Phage specificity may be especially problematic for chronic infections, which tend to be genetically diverse, suggesting they may contain variation in phage susceptibility. Such within-population variation in phage susceptibility could make phage treatment less effective, however the extent of this variation in chronic bacterial infections has not been thoroughly studied. To address this gap, we investigated variation in phage susceptibility in four populations of *Pa* derived from chronic lung infections in people with cystic fibrosis. We first measured the lytic efficiency of 12 virulent phages across 40 isolates from each patient, which revealed substantial variation in susceptibility to four LPS-targeting phages in two of four lung-derived populations. Genomic analysis suggested that within-population variation in phage susceptibility was not explained by the acquisition or loss of phage defense systems, as phage defense system carriage did not strongly vary within lung-derived populations. Conversely, phylogenetic analysis of isolates from one lung-derived population revealed that some evolutionary transitions in phage susceptibility coincided with mutations in genes involved in LPS biosynthesis. This association suggested that within-patient diversification in cell-surface features may have been responsible for variation in phage susceptibility in this population. Finally, phenotypic characterization of lung-derived isolates revealed favorable clinical associations between phage susceptibility and antibiotic resistance in one population. However, associations were largely absent in other populations, suggesting the potential to use phages to drive favorable phenotypic shifts is limited in bacterial infections with existing variation in phage susceptibility. Collectively, this work shows that phage susceptibility can vary within *Pa* chronic infections in the lungs of people with cystic fibrosis, possibly as a consequence of standing variation in cell-surface features. This variation poses a challenge for phage therapy, necessitating new approaches to ensure high efficacy in the face of extreme within-patient heterogeneity.

## Introduction

Phage therapy – the medical use of virulent bacteriophages (phages) – is an approach to control difficult-to-treat bacterial infections that has received increased interest in recent years (Kortright et al., 2019; Strathdee et al., 2023; Pirnay et al., 2024). Although phage therapy has several advantages over conventional antibiotics, such as the ability of phages to amplify at the infection site within patients, a persistent drawback is the high host specificity of individual phages (Essoh et al., 2013; Kortright et al., 2019; Hibstu et al., 2022). The efficacy of individual phages varies widely across bacterial strains, which arises from the requirement for compatibility between phages and their host cells across a complex, multi-stage infection cycle (Leprince et al., 2026). Initial phage adsorption requires compatibility between phage receptor-binding proteins (RBPs) and host cell-surface features, which often vary across bacterial strains (Harvey et al., 2018; Gaborieau et al., 2024; Müller et al., 2026; Vaitekenas et al., 2026). After genome entry, virulent (ie. obligately lytic) phages must co-opt host machinery to generate copies of their genome, package them into capsid proteins, and release these new virions via host cell lysis. Successful intracellular replication requires avoidance of or interference with phage defense systems that can prevent phage replication and assembly or cause premature host cell death before replication is completed (Labrie, Samson and Moineau, 2010). Carriage of phage defense systems varies considerably across bacterial strains, further contributing to phage host-specificity (Hussain et al., 2021; Burke et al., 2024; Costa et al., 2024). As a consequence of narrow phage host breadth, most reported cases of phage therapy involve highly personalized care where a library of phages is assessed for lytic activity against a single bacterial strain isolated from the patient, which is assumed to be representative of their infection (Chan et al., 2018; Ferry et al., 2021; 2022; Lebeaux et al., 2021; Van Nieuwenhuyse et al., 2021; 2022; Racenis et al., 2022; 2023; Cesta et al., 2023; Köhler et al., 2023; Young et al., 2023; Yang et al., 2025). Patients are then treated with one or several phages (phage cocktail) that can efficiently lyse the dominant strain responsible for their infection.

Although personalized phage therapy has been effective in many cases, treatment failure is also commonly reported (Van Nieuwenhuyse et al., 2021; Hahn et al., 2023; Aslam et al., 2024; Pirnay et al., 2024). One explanation may be that, while personalized phage therapy accounts for variation in phage susceptibility across patients, it mostly ignores the possibility of variation in phage susceptibility within single infections. Such variation may be particularly pronounced in chronic bacterial infections, which are often phenotypically and genetically heterogeneous (Mowat et al., 2011; Jorth et al., 2015; Lee et al., 2017). Within-patient variation in phage susceptibility in chronic infections could impede phage treatment in several ways. First, it could directly mitigate bacterial suppression by allowing sub-populations of phage resistant bacteria to rapidly replace any phage susceptible cells lysed during treatment. Second, standing variation in phage susceptibility could complicate phage steering – the use of phages targeting cell-surface receptors known to contribute to virulence or antimicrobial resistance in order to steer bacterial evolution towards clinically favourable traits (Chan et al., 2016; Kortright et al., 2019; Gurney et al., 2020; Ashworth et al., 2024). Phage steering assumes that *in vivo* phage resistance will evolve via loss-of-function mutations in genes encoding the phage receptor, which tend to drive predictable trade-offs *in vitro* (Gurney et al., 2020). However, if cell-surface features vary within bacterial infections prior to phage treatment, co-selection of phage resistance and clinically favourable traits may be weaker, absent, or even reversed depending on the genetic correlates of phage resistance in the population. Despite the potential for within-population heterogeneity in phage susceptibility to interfere with phage therapy, few studies have assessed variation in phage susceptibility in more than 3-5 isolates from single infections.

*Pseudomonas aeruginosa* (*Pa*) is an opportunistic bacterial pathogen that causes chronic infections in the lungs of people with cystic fibrosis (pwCF) where they are associated with decreased lung function and elevated mortality risk (Nixon et al., 2001; Courtney et al., 2007; Parkins, Somayaji and Waters, 2018). These infections are notoriously difficult to treat with conventional antibiotics, which has made them a common use case for phage therapy (Hahn et al., 2023; Pirnay et al., 2024; Chan et al., 2025). However, phage therapy for *Pa* chronic lung infections has yielded mixed results, with bacterial suppression often being short-lived and variable across patients (Hahn et al., 2023; Chan et al., 2025). The success of phage steering has also varied across patients (Chan et al., 2025). These inconsistencies in treatment outcomes may be due to the high levels of phenotypic and genetic diversity commonly observed in *Pa* chronic lung infections. Extensive within-patient diversity in *Pa* lung infections mostly result from within-patient diversification, which can be amplified by the evolution of strains with elevated mutation rates called hypermutators (Workentine et al., 2013; Feliziani et al., 2014; Jorth et al., 2015; Winstanley, O’Brien and Brockhurst, 2016; Vanderwoude et al., 2024). Isolates from the same sputum sample can vary in clinically important traits including antimicrobial resistance, motility, and the production of virulence factors like pyocyanin (Mowat et al., 2011; Workentine et al., 2013; Jorth et al., 2015; Vanderwoude et al., 2024). Additionally, substantial variation has been detected in traits related to phage infection. For example, the structure of the lipopolysaccharide (LPS), a common cell surface receptor for *Pa*-targeting phages, can vary across isolates within single infections, with O-specific antigen loss being particularly common in CF lung infections (Hancock et al., 1983; Luzar and Montie, 1985; Cullen et al., 2015; Mei et al., 2023). LPS structural variation contributed to the recent observation that *Pa* infections in the lungs of pwCF can contain substantial heterogeneity in susceptibility to R-pyocins, phage-like toxins with similar RBPs to LPS-targeting phages (Mei, Thomas and Diggle, 2021; Mei et al., 2023). Similarly, isolates from single lung infections may vary in whether they express functional type IV pili (TIVP) and/or flagella, both of which act as cell-surface receptors for *Pa*-infecting phages (Mahenthiralingam, Campbell and Speert, 1994; Workentine et al., 2013; Clark et al., 2015). These findings suggest that lung-derived populations of *Pa* may be heterogeneous in phage susceptibility, but direct tests of this heterogeneity are limited (but see Sahota et al., 2015). If within-population variation in phage susceptibility is present, it will be important to uncover its genetic determinants, both to guide sequencing-based approaches for inferring phage susceptibility and to determine whether lytic activity within single infections is mainly limited by variation in cell-surface features or phage defense systems (Burke et al., 2024; Costa et al., 2024; Müller et al., 2026; Vaitekenas et al., 2026). Further, characterizing the phenotypic correlates of phage susceptibility among isolates from single lung-derived populations will be crucial for informing whether phage steering will tend to be effective for heterogeneous chronic infections.

Here, we investigated variation in phage susceptibility in four populations of *Pa* derived from chronic lung infections in pwCF. We measured phage susceptibility across 40 *Pa* isolates from each population, revealing substantial variation in susceptibility in two lung-derived populations. Next, we examined the genetic determinants of within-population variation in phage susceptibility. These analyses focused on variation in phage defense system carriage and genetic variation in genes involved in LPS biosynthesis and phage defense or those falling within prophage regions. Finally, we asked whether phage susceptibility was associated with antibiotic resistance, biofilm production, or metrics of bacterial growth across populations. Collectively, our work is the first to characterize phage susceptibility across a large library of bacterial isolates from chronic infections, providing insight into how these heterogeneous infections may complicate phage therapy.

## Methods

### Acquiring library of lung-derived Pa isolates

The library of bacterial isolates used here was generated in a previous study exploring genetic and phenotypic diversity in lung-derived populations of *Pa* (Vanderwoude et al., 2024). Briefly, four pwCF with chronic *Pa* lung infections receiving treatment at Emory University (Atlanta, GA) provided a single sputum sample from which 75 isolates of *Pa* were collected, of which we obtained 40 (160 isolates in total). All sequencing data from this study were obtained from the NCBI SRA database (PRJNA1049984). As descried previously, each isolate was whole genome sequenced on an Illumina NovaSeq, which produced 250 bp paired-end reads at a mean coverage of 70-fold (Vanderwoude et al., 2024). In addition, one random isolate was selected as a reference strain for each population and received Oxford Nanopore Technology (ONT) sequencing to facilitate genome assembly (Vanderwoude et al., 2024). Additionally, the susceptibility of the full *Pa* isolate collection to six chemical antibiotics (amikacin, meropenem, piperacilin-tazobactam, ciprofloxacin, tobramycin, and ceftazidime) was measured using disc diffusion assays in the prior study (Vanderwoude et al., 2024).

### Acquiring phage panel

We selected a panel of 12 virulent phages from a larger library of phages assembled by the Yale Center for Phage Biology and Therapy (YCPBT). Phages were only considered if they could amplify to a high titer (> 10^9^ PFU/mL) on the lab strain MPAO1. Phages were selected to maximum the lytic breadth of the panel on a library of clinical isolates also collected by the YCPBT based on spot assays (unpublished data).

To determine the cell-surface receptor of each phage in our panel, high titer stocks of each phage were plated in a 10-fold dilution series in 10 µL spots on lawns of MPAO1 as well as a set of MPAO1-derived knock-out strains. Knock-out strains included strains lacking the TIVP as well as various components of the LPS. All knock-out strains were created in a genetic background lacking both filamentous prophages normally present in the MPAO1 genome (Pf4 and Pf6), which we refer to as MPAO1ΔΔ. Lawns were generated by first growing each bacterial strain overnight (16 h) at 37 °C in 1.5 mL of lysogeny broth (LB) in 24-well plates. 200 µL of each overnight culture was then added to 4 mL of LB top agar (0.75% w/v agar) and spread onto LB agar plates (1.5% w/v agar). The efficiency of plating (EOP) of phages across the knock-outs was quantified by computing the quotient of the inferred density of each phage on each knock-out relative to the reference strain (MPAO1). EOP was measured in duplicate.

### Quantifying phage susceptibility

The susceptibility of each of the 160 isolates in our bacterial library to each of the 12 phages in our panel was measured via EOP assay, similar to the assay described above. Due to the relatively slow growth rate of some isolates in our library, bacteria were grown for 24 h rather than overnight. Bacterial lawns of each clinical isolate as well as MPAO1 were generated as described above. Phage titers were adjusted to 10^9^ PFU/mL and were then plated on bacterial lawns in 2 µL spots at dilution factors ranging from 10^1^-10^6^. EOP was measured in duplicate.

### Analysis of bacterial genomes

To generated high-quality annotations of the four reference strains from each population, ONT reads from these strains were first filtered for quality using *fastplong*. Subsequently, assemblies of each reference strain were generated using *flye* and were then polished using *medaka* and *polypolish*, the latter of which used short-read sequencing data from the same isolates. Assemblies were then annotated using *bakta* with a custom library of protein sequences acquired from three *Pseudomonas spp.* protein libraries available on UniProt: PAO1 (taxon ID: 208964), PA14 (taxon ID: 208963), and PA7 (taxon ID: 381754). This generated high-quality reference genome annotations which were used for subsequent bioinformatic steps.

The four reference genomes were further analyzed to determine loci of predicted prophages as well as genes predicted to be involved in phage defense. Intact prophage regions were predicted using *phastest* and were subsequently used to determine which genetic variants in each population fell within predicted prophages. Defense genes in each reference genome were predicted using both *padloc* and *defense-finder*. Duplicate phage defense genes detected by both programs were removed. These outputs were used to determine which genetic variants fell into predicted phage defense genes.

Genetic variants present in each population were detected by comparing short-read sequences from each isolate to the corresponding within-population reference genome. First, the short-read sequences were filtered for quality using *fastp*. Then, *breseq* was used to call variants in each isolate and generate a multi-sequence alignment of variable sites in each population. This alignment was used to generate a maximum likelihood phylogeny with *IQtree* using standard parameters. Phylogenies for each population were visualized in R using the package *ggtree*. Given the large number of genetic variants in some of our populations (Vanderwoude et al., 2024), only variants falling into three categories were considered in detail: those occurring in genes involved in LPS biosynthesis, in genes falling in predicted prophage regions, and in predicted phage defense genes. The list of genes involved in LPS biosynthesis was generated by first downloading a full set of genes found in the three *Pseudomonas spp.* reference strains mentioned above and subsetting them for genes with gene ontology (GO) terms related to LPS (see Supplementary Materials for list of GO terms used). This list was supplemented with a comprehensive set of LPS-related genes in *Pa* provided in King et al. (2009). Genetic variants falling in predicted prophage regions or phage defense genes were defined by comparing the loci of the variants to prophage and phage defense loci determined above.

### Bacterial growth curves

Microplate growth curves were run for each isolate in our collection to estimate their growth performance in the absence of added phages. Growth curves used the same 24-h cultures used in the EOP assay. Bacterial cultures were first diluted 1:10 in LB before 20 µL of these pre-diluted cultures was added to 180 µL of fresh LB in a well of a 96-well plate. Plates were incubated at 37 °C under constant double-orbital shaking for 24 h and optical density (600 nm) was measured every 10 minutes. The R package *gcplyr* was used to compute the maximum per capita growth rate, maximum density, and area-under-the-curve for each growth curve (Blazanin, 2024). Growth curves for bacterial isolate were performed in duplicate.

### Quantifying biofilm formation

Biofilm formation in each strain in the bacterial isolate library was quantified using a microplate biofilm formation assay (O’Toole, 2011). Briefly, each isolate was grown overnight in LB and diluted 1:100 into M63 minimal media supplemented with magnesium sulfate (1 mM), glucose (0.2% w/v), and casamino acids (0.5% w/v). 100 µL of the diluted culture was then transferred into a well of a 96-well plate and incubated for 24 h at 37 °C. After incubation, the cultures were dumped out, and the plate was rinsed in water twice. Then, 125 µL of crystal violet solution (0.1% w/v) was added to each well and allowed to incubate at room temperature for 15 min. Then, plates were rinse three times with water to remove excess stain and allowed to dry overnight. The next day, 125 µL of 30% v/v acetic acid was added to each well and allowed to incubate at room temperature for 15 min to solubilize the stain. Finally, the volume was transferred to a fresh 96-well plate and the absorbance (550 nm) was measured to quantify the biofilm formation. All isolates were measured in triplicate. 30% v/v acetic acid was used as a negative control.

### Statistics

Statistical analyses were performed using R (R Core Team, 2026). The relationships between phage susceptibility and clinically relevant bacterial phenotypes were assessed using individual linear models for each combination of phage (n = 4), phenotype (n = 10), and population (n = 4) for a total of 160 models. For each model, log_10_ EOP was used as a continuous predictor, and all response phenotypes were also continuous. The Benjamini-Hochberg method was used to adjust P-values to control for multiple comparisons.

## Results

### Panel of 12 MPAO1-infecting phages includes TIVP- and LPS-targeting phages

We acquired a panel of 12 phages from the Yale Center for Phage Biology and Therapy that had the maximum collective host breadth on an in-house collection of clinical *P. aeruginosa* isolates (data not shown). We then used a set of genetic knock-outs to determine the host receptor of each of the 12 phages. 7/12 phages but had no detectable plaquing on mutants with disrupted type IV pili (MPAO1ΔΔΔ*pilA*), suggesting that these 7 phages (31BULL2, Φ5, 8BT, PB, 25S2, 40S1, and 34T) are TIVP-dependent (Figure 1). Each of the remaining 5 phages (LPS-5, Φ45, Φ6, INF, and Φ10) showed reduced plaquing efficiency on some combination of LPS biosynthesis knock-outs (Figure 1). This implied that these 5 phages are LPS-dependent. Although the specific LPS residues targeted by these phages could not be determined, the moderate-to-high efficiency with which all LPS-dependent phages plaqued on the Δ*wzm*Δ*wzy* and Δ*wbpL* strains suggests that none of them require either the O-specific antigen or the common polysaccharide antigen to infect MPAO1 (Figure 1).

**Figure 1:**
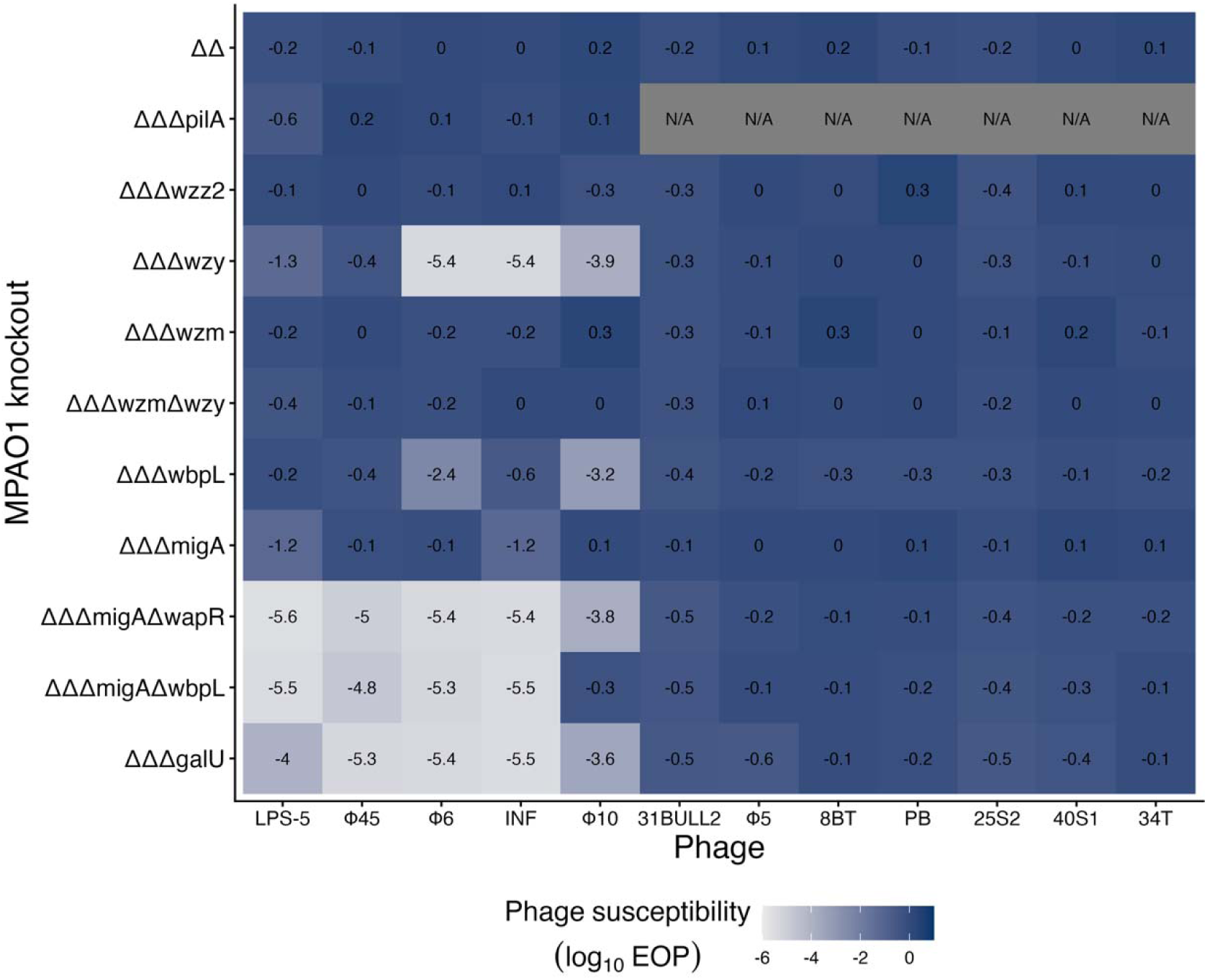
Susceptibility of TIVP, LPS knock-outs to panel of 12 virulent bacteriophages. Tiles represent mean phage susceptibility (n = 2) relative to MPAO1. All knockouts were generated in MPAO1ΔPf4ΔPf6 background (abbreviated to “ΔΔ” here). “N/A” values indicate phage susceptibility was below the limit of detection.

### Phage susceptibility varies within and among lung-derived populations of Pa

We acquired 40 isolates of *Pa* collected from each of four sputum samples taken from CF patients with chronic lung infections at a single timepoint. The clinical histories of each patient are described in Vanderwoude et al. (2024). Patients were all infected with different single strains of *Pa* but within-population genetic diversity varied considerably across patients (Vanderwoude et al., 2024).

We measured the susceptibility of the full isolate library to our panel of 12 phages using efficiency of plating (EOP). We found that 8 phages (Φ10 as well as all of the TIVP-targeting phages) produced plaques on only 1-2 isolates across the entire bacterial library (Supplementary Figure 1). Conversely, the remaining four LPS-targeting phages had relatively high host breadth in the collection, each forming plaques on at least 52 isolates (range: 52-103). We refer to these phages as “broad host-range phages” below.

Consistent with expectations, we found that phage susceptibility substantially varied within 2/4 lung-derived populations (Figure 2). In CF1, we detected susceptibility to one of our four broad host-range phages in 28-50% of isolates and susceptible isolates ranged in log_10_ EOP from ∼0 (equal EOP to MPAO1) to ∼-4 (10,000-fold reduction in phage efficiency; Figure 1A). In CF3, we detected susceptibility to broad host-range phages in 0-93% of isolates and susceptible isolates similarly varied from highly susceptible (log_10_ EOP ∼0) to extremely weakly susceptible (log_10_ EOP ∼-5; Figure 1C). Conversely, phage susceptibility was less variable in Populations CF2 and CF4, which were largely phage resistant (0-15% susceptible) and universally phage susceptible, respectively (Figure 1BD). These results suggested that lung-associated populations of *Pa* can contain significant heterogeneity in phage susceptibility, while other populations contain little detectable variation in phage susceptibility.

**Figure 2:**
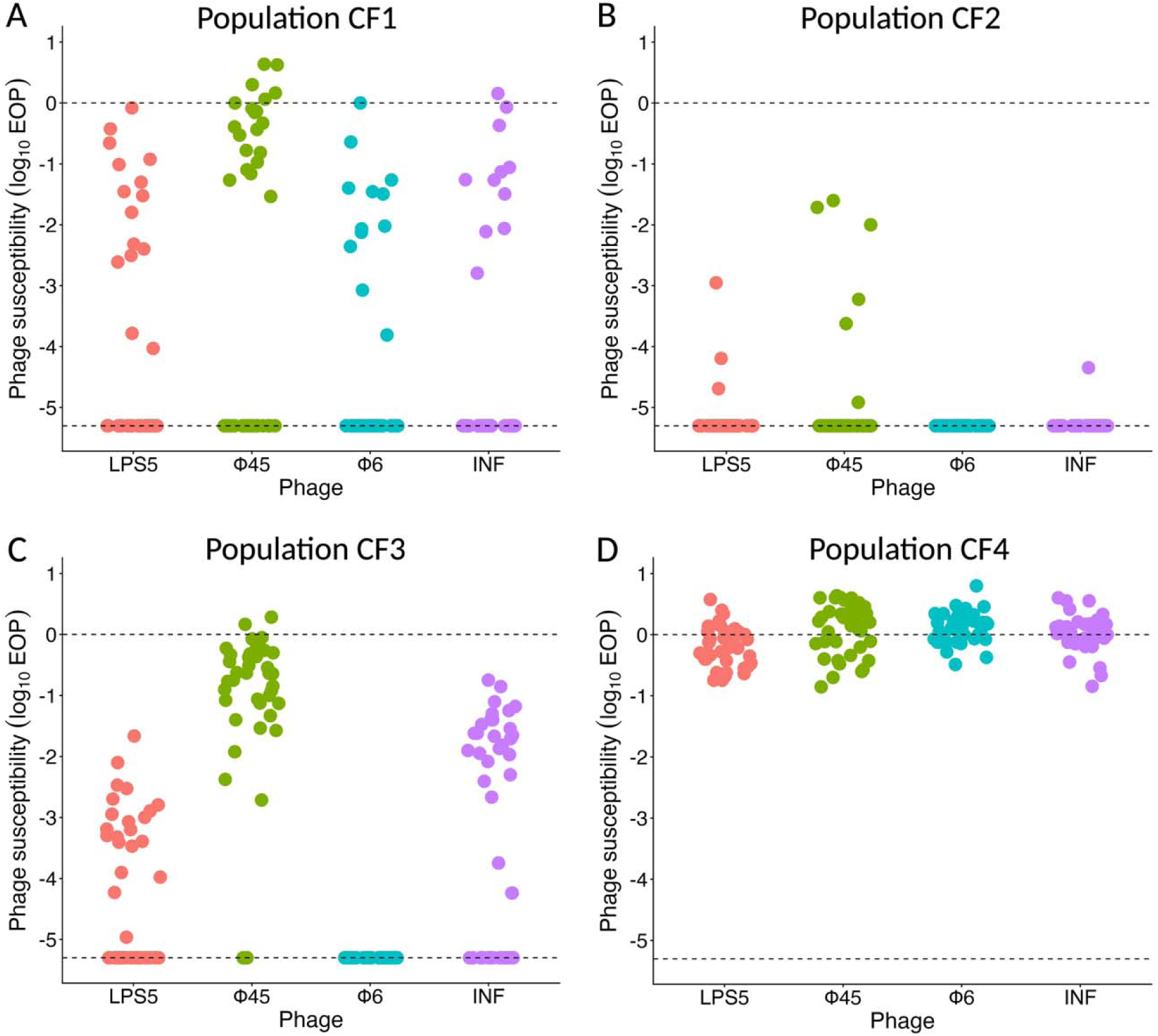
Susceptibility of library CF lung-derived isolates of *Pseudomonas aeruginosa* to four LPS-targeting virulent phages. Each point represents the mean susceptibility (n = 2) of a single bacterial isolate from (A) Population CF1, (B) Population CF2, (C) Population CF3, and (D) Population CF4 to one of four LPS-targeting phages. Lower dashed lines represent the lower limit of detection (log_10_ EOP = −5.3) and upper dashed lines represent equal susceptibility of clinical bacterial isolates to the reference strain, MPAO1.

### Defense system carriage varies little within lung-derived populations, varies substantially across populations

To investigate the genetic determinants of variation in phage susceptibility within lung-derived populations, we first annotated the phage defense systems present in all 160 isolates in the collection. This revealed that defense system presence was largely invariant within populations, with only one defense system in one population falling between 7.5-97.5% frequency (PDC-M11 in Population CF1; Figure 3). Conversely, defense system carriage varied substantially across populations, with only 6/37 defense systems found in more than one population (Figure 3). Populations also varied in the number of defense systems carried, ranging from 10 systems in CF1 and CF2 to 17 systems in CF3 (Figure 3). Collectively, these results suggested that variation in defense system carriage did not explain variation in phage susceptibility within our lung-derived populations of *Pa*.

**Figure 3:**
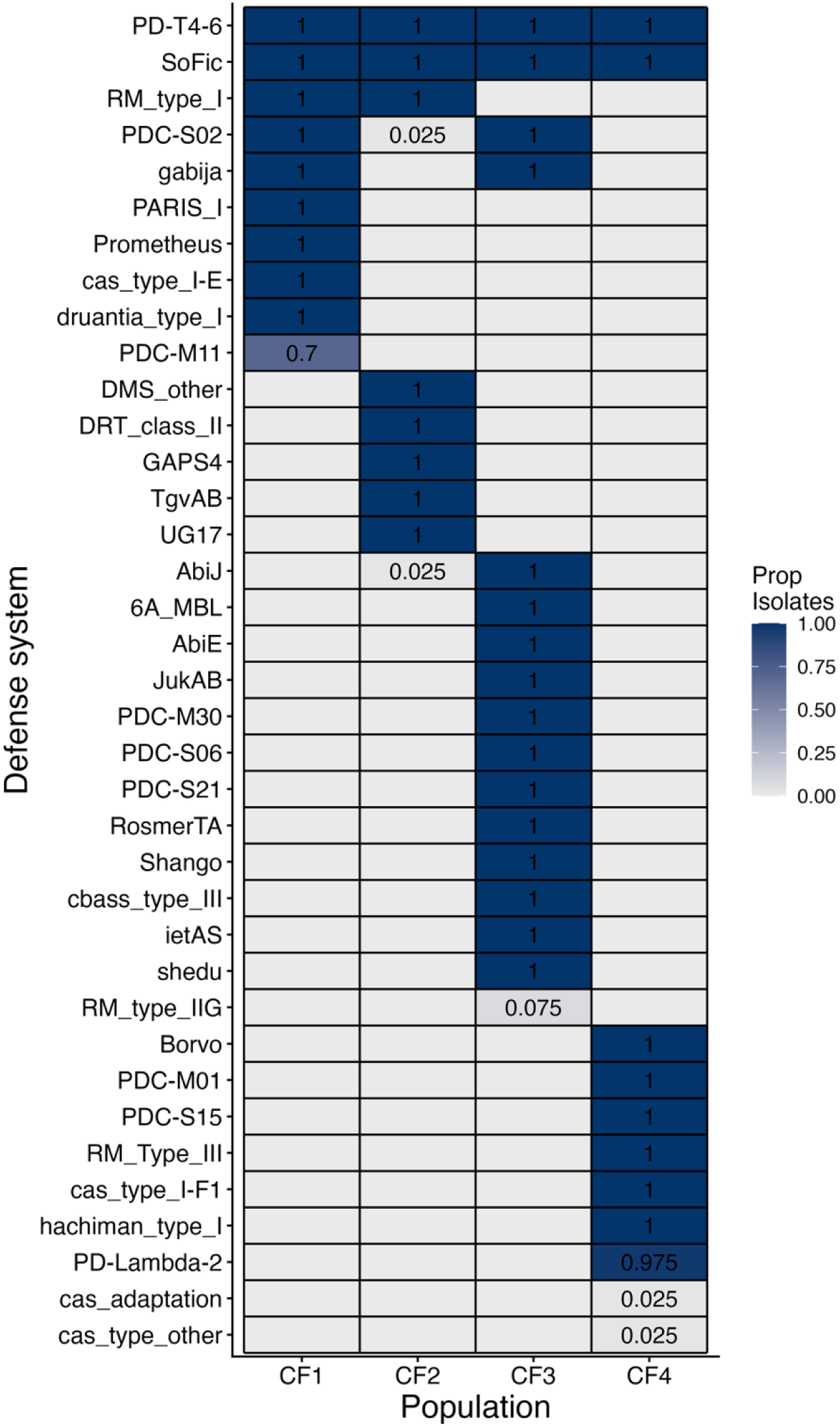
Defense system carriage across four CF lung-derived populations of *P. aeruginosa*. Defense systems were annotated in 40 isolates from each of the four CF-lung derived populations using both *DefenseFinder* and *Padloc*. The proportion of isolates in each population carrying each defense system is represented by the color of each tile, as well as numerically within each tile when the proportion exceeded 0.

### Phylogenetic transitions in phage susceptibility coincident with mutations in LPS-related genes in CF1

We generated a maximum-likelihood phylogeny of all isolates from Population CF1 and annotated mutations in LPS-related genes, genes in prophage regions, and phage defense genes along their branches (Figure 4). This revealed that two evolutionary transitions in phage susceptibility in Population CF1 were associated with mutations in LPS-related genes (Figure 4). Specifically, the emergence of susceptibility in the branch leading to Clade 2 (medium grey) co-occurred with two mutations in LPS-related genes: *hldE* and *waaP* (Figure 4). Both of these genes are involved in LPS inner core biosynthesis (King et al., 2009). In addition, the emergence of susceptibility in the branch leading to Clade 1 (light grey) co-occurred with three other mutations in LPS-related genes: *msbA* and *lpxO2*, which are involved in lipid A biosynthesis, and *waaL*, the O-antigen ligase (King et al., 2009). This branch also contained a mutation in *cas3e*, a key component of the CRISPR Type I-E phage defense system, as well as two mutations in a prophage region unrelated to LPS or phage defense (Westra et al., 2012). In contrast to these examples, the emergence of phage susceptibility in Clade 3 was not associated with mutations in any LPS, prophage, or phage defense gene, suggesting other factors were responsible for phage susceptibility in these isolates. Collectively, these results are consistent with within-patient genetic diversification, particularly in LPS-related genes, driving the variation in phage susceptibility we observed in Population CF1.

**Figure 4:**
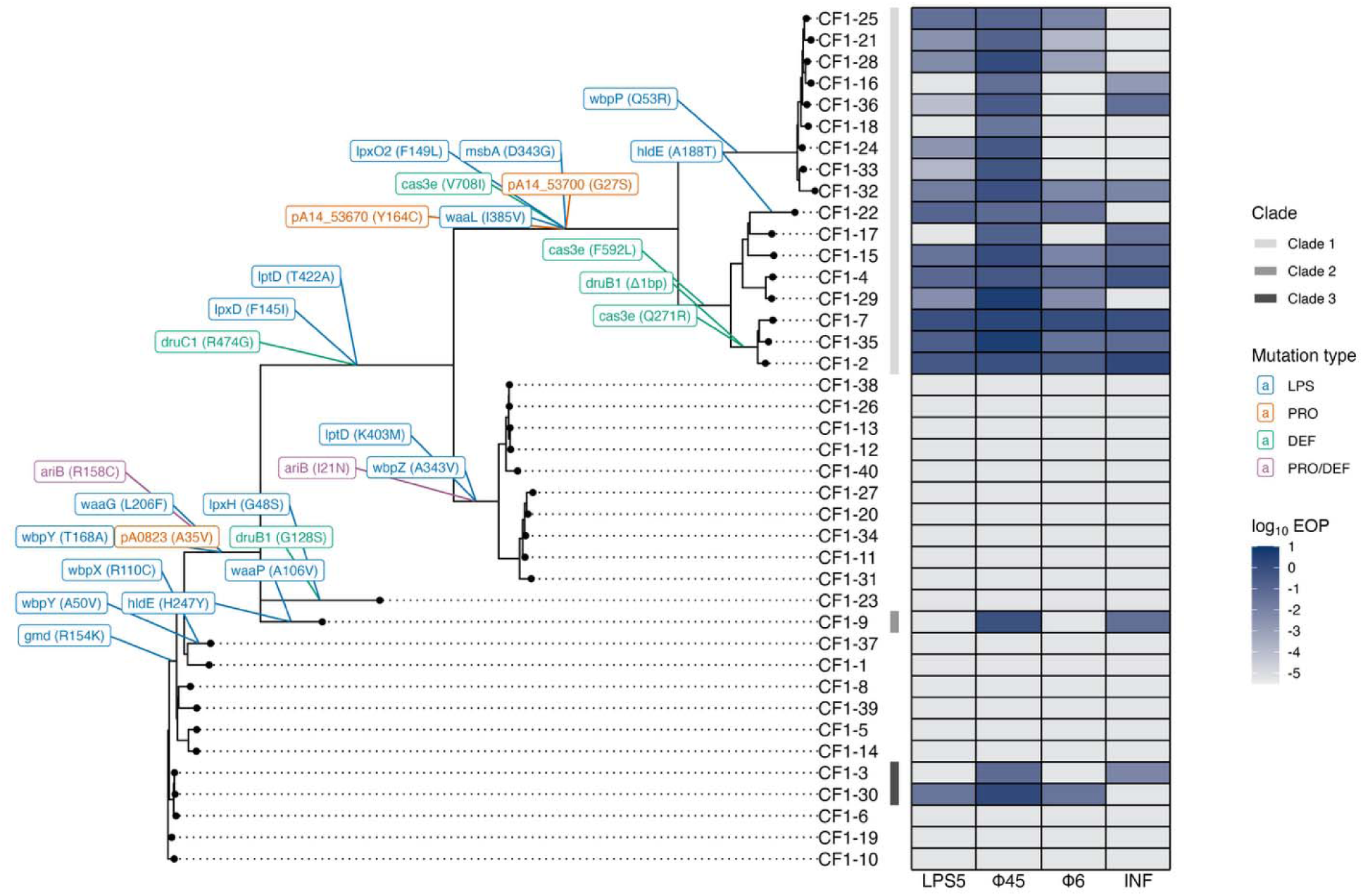
Evolutionary genetics of phage susceptibility in lung-derived Population CF1. Maximum-likelihood phylogeny of isolates from Population CF1 (left) with EOP values across four broad host range phages provided in a tile plot along the tips (right). Non-synonymous SNPs and small indel occurring in coding regions of LPS genes (LPS), prophage regions (PRO), and phage defense genes (DEF) are annotated along branches of the phylogeny. Mutations in phage defense genes found within prophage regions annotated as PRO/DEF mutations. Clades containing phage susceptible isolates are highlighted with vertical bars adjacent to tips.

### Associations between phage susceptibility and clinically important traits are variable across populations

To assess the potential for phage steering in heterogeneous chronic *Pa* lung infections, we asked whether increased phage susceptibility was associated with clinically unfavourable traits within our lung-derived populations (Figure 5). We found several clinically desirable associations in Population CF1, such as the positive association between phage susceptibility and biofilm production and the negative associations between phage susceptibility and TZP, CAZ, and AK sensitivity (Figure 5A; *P* < 0.05). Conversely, significant associations between phage susceptibility and clinically important traits were virtually absent in the other three populations (Figure 5; Supplementary Figure 2). One exception was the negative association between Φ45 susceptibility and biofilm production in CF3, which is clinically undesirable (phage addition expected to drive increased biofilm production). Overall, these results indicate that clinically favorable associations between phage susceptibility and other bacterial traits may exist in some lung-derived populations of *Pa*, but they are not universal and may occasionally be reversed.

**Figure 5:**
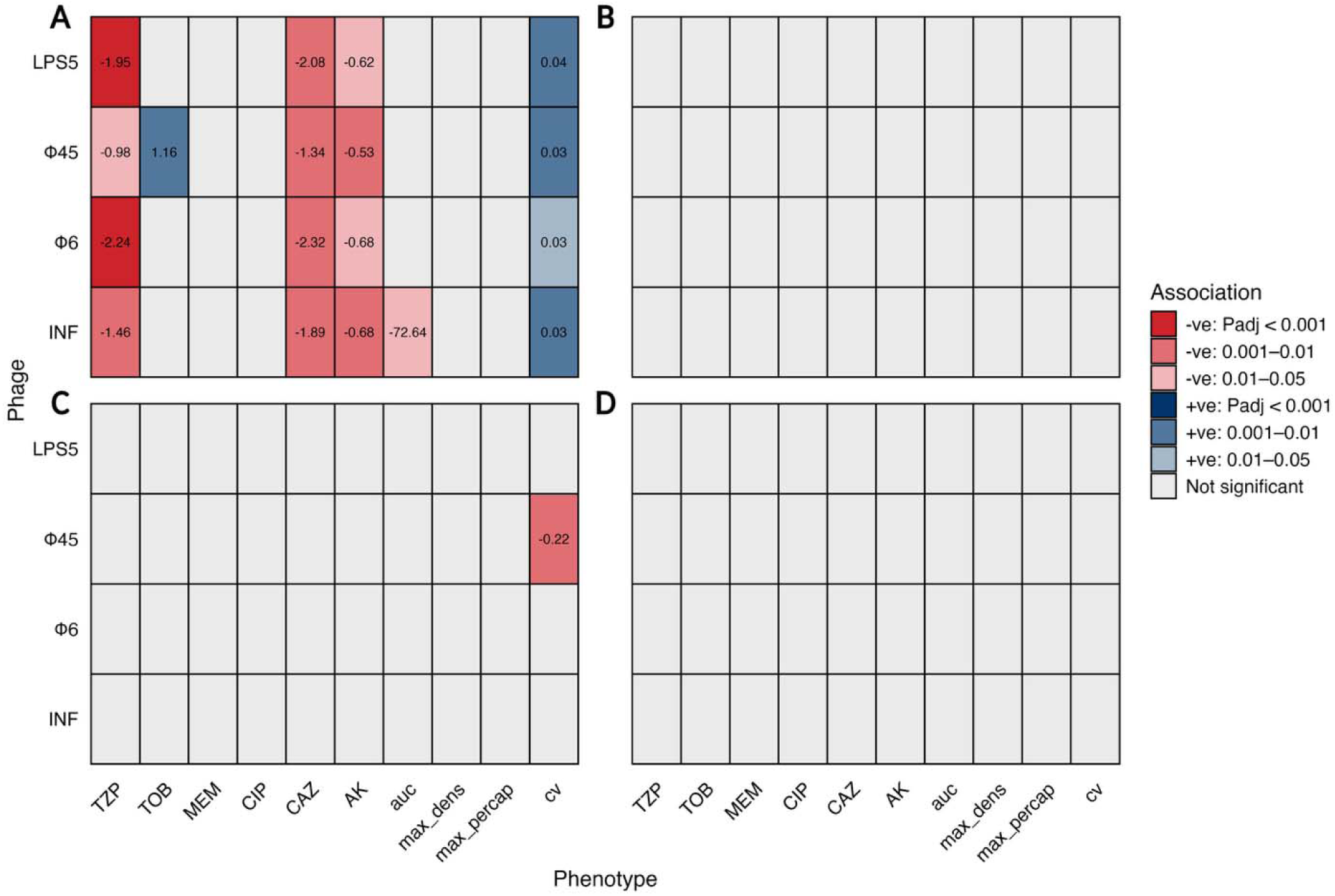
Relationship between phage susceptibility and clinically important phenotypes across four lung-derived populations of *P. aeruginosa*. Each tile represents the linear association between susceptibility to broad host range phages (log_10_ EOP) and clinically important phenotypes, including antibiotic susceptibility (mm), area-under-the curve (auc; OD_600_/min), maximum density (max_dens; OD_600_), and maximum per capita growth rate (max_percap; min^−1^) from microplate growth curves, as well as biofilm formation (cv; OD_550_). Associations shown for (**A**) Population CF1, (**B**) Population CF2, (**C**) Population CF3, and (**D**) Population CF4. Tile color represents the statistical significance of relationships after correcting for multiple comparisons, with grey tiles representing non-significant relationships. For significant relationships, slopes are shown inside tiles.

## Discussion

Within-population variation in phage susceptibility could decrease the efficacy of phage therapy for chronic bacterial infections. However, few studies have assessed phage susceptibility in more than 3-5 isolates from single infections. Here, using large bacterial isolate libraries collected from single infections (40 isolates/patient), we showed that populations of *P. aeruginosa* derived from the lungs of pwCF can contain substantial variation in phage susceptibility. Genomic analyses suggested that within-population variation in phage susceptibility may have been driven by within-patient diversification in cell-surface features rather than variation in defense system carriage among isolates from the same population. Additionally, we found clinically favorable statistical associations between phage susceptibility and antibiotic susceptibility and biofilm formation in one population but not the other three. This latter result suggests that the efficacy of phage steering may be variable in heterogenous chronic bacterial infections.

Over the last 15 years, the challenges posed by phenotypic and genetic heterogeneity for the management of *Pa* chronic lung infections have received increased attention (Mowat et al., 2011; Workentine et al., 2013; Clark et al., 2015; Darch et al., 2015; Jorth et al., 2015; Winstanley, O’Brien and Brockhurst, 2016; Mei, Thomas and Diggle, 2021; Vanderwoude et al., 2024). *Pa* isolates from these infections often vary in antibiotic sensitivity, biofilm production, and virulence both within and across clinical samples (Mowat et al., 2011; Workentine et al., 2013; Jorth et al., 2015; Vanderwoude et al., 2024). Our work adds to this picture by demonstrating that coexisting genotypes can also vary in susceptibility to virulent phages (Figure 2). Our results contrast somewhat with Sahota et al. (2015), which found that several phages lysed over 95% of a library of 80 lung-derived strains of *Pa* derived from two patients with chronic lung infections. The difference in observed heterogeneity between these studies may reflect that some proportion of *Pa* chronic lung infections contain substantial variation in phage susceptibility while others are relatively homogeneous. When heterogeneity in phage susceptibility is present, it may cause similar problems for phage therapy as it does for more antibiotic therapy. Specifically, heterogeneity would both complicate pre-treatment phage susceptibility testing and allow phage resistant sub-populations to rapidly outgrow phage susceptible strains during phage treatment. If so, our results may explain why phage treatment for chronic *Pa* lung infections has had variable outcomes (Hahn et al., 2023; Chan et al., 2025). Given the increased reliance on phage therapy to control chronic *Pa* lung infections, understanding the relationship between within-population variation in phage susceptibility and phage therapy outcomes should be a priority for future work.

Whether variation in bacterial cell-surface features or phage defense systems are more responsible for phage host specificity has been the subject of recent debate (Burke et al., 2024; Costa et al., 2024; Leprince et al., 2026; Müller et al., 2026; Vaitekenas et al., 2026). Currently evidence in *Pa* is mixed, suggesting that both factors are important in shaping phage host range across diverse strains of *Pa* (Burke et al., 2024; Müller et al., 2026; Vaitekenas et al., 2026). Our work is consistent with cell-surface features being most important for explaining variation in phage susceptibility among closely related strains from the same population. Specifically, we found that phage defense system carriage did not vary substantially in single populations, making it unlikely to explain within-population variation in phage susceptibility (Figure 3). Conversely, focusing on Population CF1, we found that mutations in LPS biosynthesis genes were co-incident with two evolutionary transitions in phage susceptibility (Figure 4). In particular, our observation that a mutation in *waaL*, which encodes the O-antigen ligase in *Pa*, was associated with the emergence of a highly phage susceptible clade is notable (Figure 4, Clade 1). *Pa* populations often lose the O-specific antigen of LPS during pathoadaptation, which can occur via *waaL*-inactivation (Hancock et al., 1983; Luzar and Montie, 1985; Abeyrathne et al., 2005; Cullen et al., 2015). Given the importance of the O-antigen for phage adsorption, the *waaL* mutation we detected is a strong candidate for explaining the transition from phage resistance to phage susceptibility in Clade 1. However, Population CF1 contains over 1,500 SNPs and small indels (Vanderwoude et al., 2024), which limits our ability to infer which mutation(s) were ultimately responsible for these evolutionary transitions. Future work will evaluate the impact of a set of candidate phage susceptibility-conferring mutations, including the observed *waaL* mutation, on phage susceptibility. If within-population variation in phage susceptibility is primarily explained by variation in host cell-surface features, this would add support for phage engineering efforts aimed at diversifying the RBPs of virulent phages (Yehl et al., 2019; Lenneman et al., 2021; Sikkema et al., 2026).

One of the commonly cited advantages of phage therapy is the ability to leverage trade-offs between phage resistance and clinically harmful traits like antibiotic resistance and biofilm production (Kortright et al., 2019; Gurney et al., 2020). This approach, called phage steering, relies on genetic correlations between phage resistance and clinically benign traits. In lab experiments using genetically homogeneous bacterial populations, phage mediated selection often drives predictable trade-offs, most likely due to harsh phenotypic costs associated with *de novo* phage receptor loss for bacteria (German and Misra, 2001; León and Bastías, 2015; Chan et al., 2016; Burmeister et al., 2020; Kortright et al., 2022; Chen et al., 2024; Houpt, Hernandez and Turner, 2026). For instance, selection from phages targeting the LPS, which impacts the permeability of cell membranes to antibiotics, has been shown to drive increased colistin susceptibility in *Escherichia coli* (Burmeister et al., 2020). However, in chronic *Pa* lung infections, phage steering has yielded mixed success, with phage treatment failing to drive expected trade-offs in some cases (Chan et al., 2025). Our results provide a potential explanation for phage steering failure: variable genetic correlations between phage resistance and clinically benign traits across infections (Figure 5). These results raise the question of why genetic correlations vary across populations. One possible reason is that evolution within chronic infections, which can persist for decades, may allow for compensatory evolution following diversification in cell-surface features, similar to that which follows the evolution of costly antibiotic resistance (Levin, Perrot and Walker, 2000; Melnyk et al., 2017). Compensatory evolution could mitigate the pleiotropic consequences of mutations in cell-surface features, making them less likely to be associated with clinically important traits like antibiotic resistance. Although more work into the phenotypic correlates of phage susceptibility in chronic bacterial infections is clearly needed, our results are not promising for the generalizability of phage steering for chronic bacterial infections. One caveat is the limited cell surface receptor diversity of our broad host range phages, which all targeted the LPS (Figure 1). It is possible that, for example, we would have detected more consistent associations between phage resistance and antibiotic sensitivity if we used phages targeting proteins involved in antibiotic efflux such as OprM (Chan et al., 2016).

Phage therapy represents a much-needed new approach in western medicine for controlling antibiotic resistant bacterial infections. Despite its many high-profile successes (Schooley et al., 2017; Chan et al., 2018; 2025; Pirnay et al., 2024), more work is needed to address the limitations of phage therapy to improve its overall efficacy and breadth of use. Here, we highlighted how narrow phage host ranges could limit the efficacy of phage therapy for heterogeneous chronic bacterial infections. To address this limitation, approaches to clinical testing and phage selection may need to be adjusted to account for within patient heterogeneity. In particular, we view the potential for phage modification, whether via phage training or genetic engineering, as a promising approach to mitigate the challenges posed by genetically heterogenous chronic infections (Borin et al., 2021; Lenneman et al., 2021; Ngiam, Weynberg and Guo, 2024).

## Supporting information

King LPS Genes

LPS GO Accessions

## Acknowledgments

We thank members of the Turner Lab, especially Benjamin K. Chan, Kaitlyn E. Kortright, Michael Blazanin, Catherine A. Hernandez, and Albert C. Vill for their helpful advice and technical assistance on this project. We acknowledge support from Yale University for the Center for Phage Biology & Therapy and from the Yale Institute for Biospheric Studies, which funded this work. NSBH was also supported by a National Sciences and Engineering Research Council of Canada Postgraduate Scholarship (CGSD3 - 559651 - 2021).

