## Supplementary Figures for "Bacteriophage susceptibility varies within chronic *Pseudomonas aeruginosa* lung infections"

**Short Title**: Within-patient variation in phage susceptibility

**Authors**: Noah S. B. Houpt^1,*^, Jelly Vanderwoude^2^, Stephen P. Diggle^2^, Paul E. Turner^3,4,5^

**Keywords**: Phage therapy, genetic variation, phenotypic heterogeneity, trade-offs, lipopolysaccharide, phylogeny, cystic fibrosis, phage defense

^1^Department of Ecology & Evolutionary Biology, Yale University, New Haven, CT, USA

^2^Center for Microbial Dynamics and Infection, School of Biological Sciences, Georgia Institute of Technology, Atlanta, GA, USA

^3^Center for Phage Biology and Therapy, Yale University, New Haven, CT, USA

^4^Yale Quantitative Biology Institute, Yale University, New Haven, CT, USA

^5^Program in Microbiology, Yale School of Medicine, New Haven, CT, USA

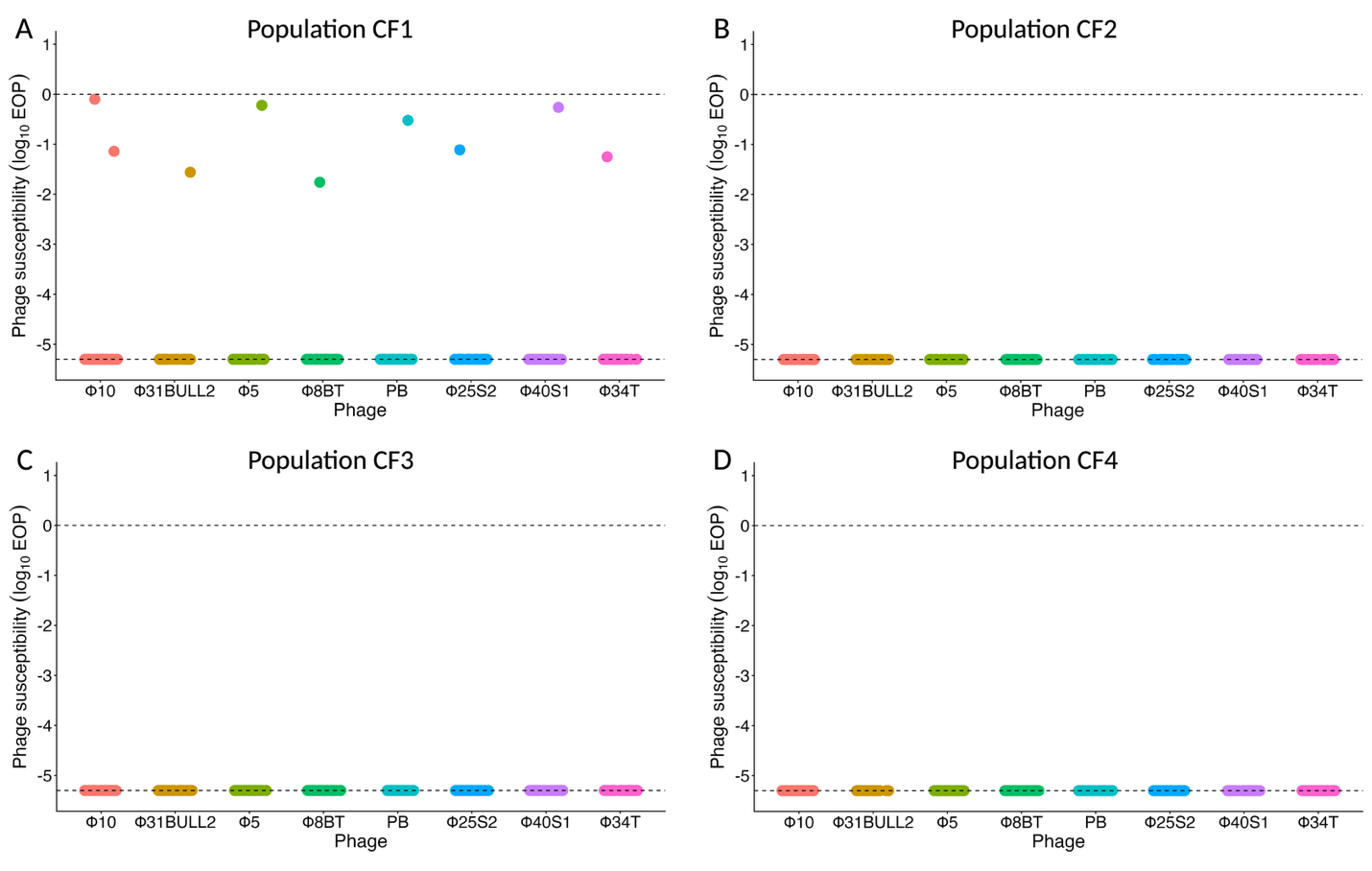

**Supplementary Figure 1:** **Susceptibility of library CF lung-derived isolates of *Pseudomonas aeruginosa* to 8 narrow host range phages**. Each point represents the mean susceptibility (n = 2) of a single bacterial isolate from (A) Population CF1, (B) Population CF2, (C) Population CF3, and (D) Population CF4 to one of four LPS-targeting phages. Lower dashed lines represent the lower limit of detection (log_10_ EOP = -5.3) and upper dashed lines represent equal susceptibility of clinical bacterial isolates to the reference strain, MPAO1.

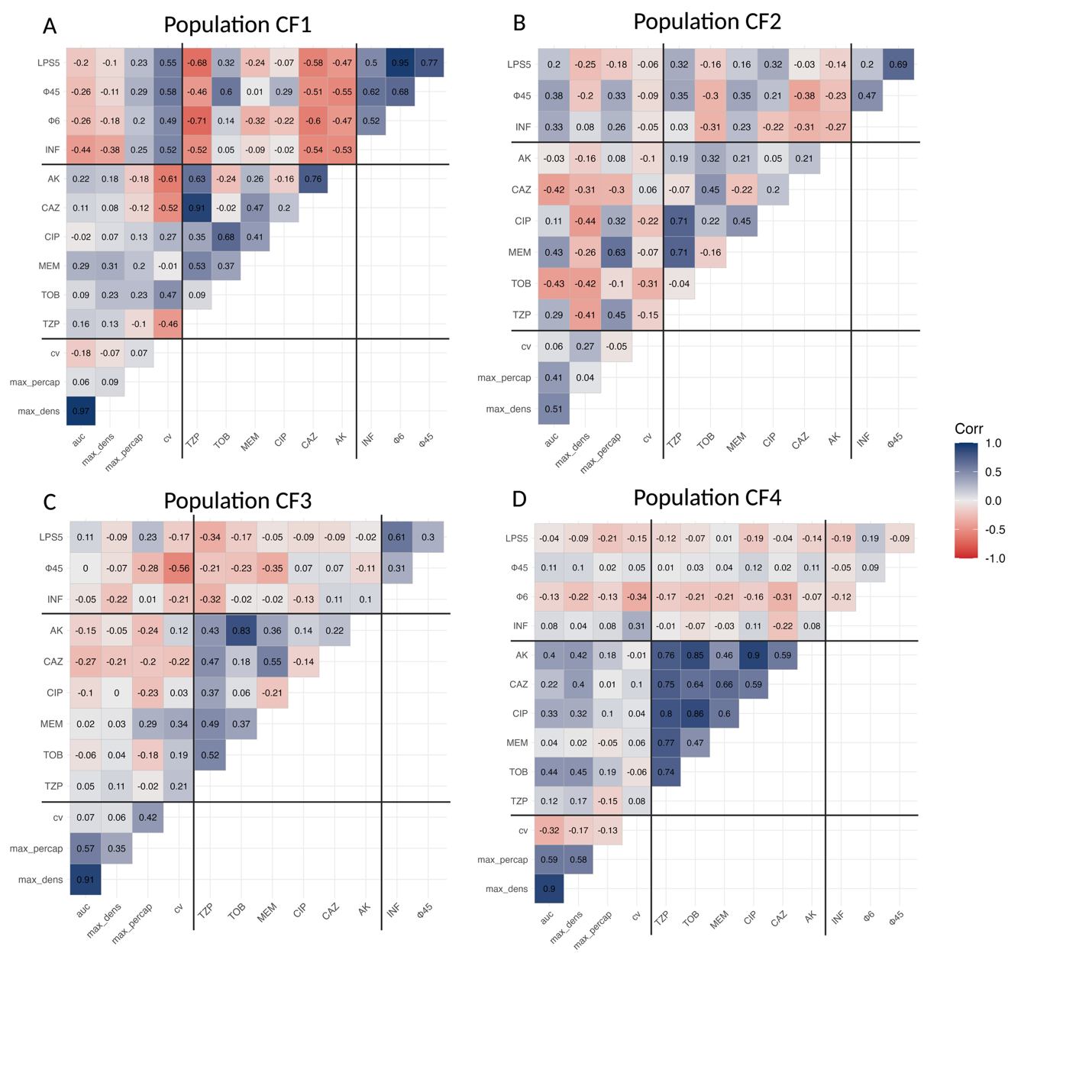

**Supplementary Figure 2: Correlation between clinically important traits in four lung-derived populations of *Pseudomonas aeruginosa*.** Tiles display the Spearman’s correlation coefficient between all combinations of susceptibility to broad host range phages (log_10_ EOP), antibiotic susceptibility (mm), area-under-the curve (auc; OD_600_/min), maximum density (OD_600_), and maximum per capita growth rate (min^-1^) from microplate growth curves, as well as biofilm formation (cv; OD_550_). Black lines demarcate groups of phenotypes (phage susceptibility, antibiotic susceptibility, bacterial growth traits and biofilm production).
